# Characterization and function of medium and large extracellular vesicles from plasma and urine by surface antigens and Annexin V

**DOI:** 10.1101/623553

**Authors:** Ko Igami, Takeshi Uchiumi, Saori Ueda, Kazuyuki Kamioka, Daiki Setoyama, Kazuhito Gotoh, Masaru Akimoto, Shinya Matsumoto, Dongchon Kang

## Abstract

Medium/large extracellular vesicles (m/lEVs) are released by most cell types and are involved in multiple basic biological processes. Analysis of m/lEV levels in blood or urine may help unravel pathophysiological findings in many diseases. However, it remains unclear how many naturally-occurring m/lEV subtypes exist as well as how their characteristics and functions differ from one another. Here, we identified m/lEVs pelleted from plasma and urine samples by differential centrifugation and showed by flow cytometry that they typically possessed diameters between 200 nm and 800 nm. Using proteomic profiling, we identified several proteins involved in m/lEV biogenesis including adhesion molecules, peptidases and exocytosis regulatory proteins. In healthy human plasma, we could distinguish m/lEVs derived from platelets, erythrocytes, monocytes/macrophages, T and B cells, and vascular endothelial cells using various surface antigens. m/lEVs derived from erythrocytes and monocytes were Annexin V positive. In urine, 50% of m/lEVs were Annexin V negative but contained various membrane peptidases derived from renal tubular villi. Urinary m/lEVs, but not plasma m/lEVs, showed peptidase activity. The method we have developed to characterize cell-derived m/lEVs suggests the possibility of clinical applications.

## Introduction

Extracellular exosomes and microvesicles play essential roles in cell-cell communication and are diagnostically significant molecules. Extracellular vesicles (EVs) are secreted from most cell types under normal and pathophysiological conditions [1, 2]. These membrane vesicles can be detected in many human body fluids (blood, urine, cerebrospinal fluid, bile, semen, synovial fluid, saliva, breast milk and malignant ascites) and are thought to have signaling functions in interactions between cells. Analysis of EVs may have applications in therapy, prognosis, and biomarker development in various fields. The hope is that using EV analysis, clinicians will be able to detect the presence of disease as well as to classify its progression using noninvasive methods such as liquid biopsy [3, 4].

Medium/large extracellular vesicles (m/lEVs) can be classified based on their cellular origins, biological functions and biogenesis [5]. In a broad sense, they can be classified into m/lEVs with diameters of 100-1000 nm diameter (membrane blebs) and smaller EVs (e.g. exosomes) with diameters of 30-150 nm [6, 7]. The m/lEVs are generated by direct outward budding from the plasma membrane [8], while smaller EVs (e.g. exosomes) are produced via the endosomal pathway with formation of intraluminal vesicles by inward budding of multivesicular bodies (MVBs) [7]. MVBs subsequently fuse with the plasma membrane, releasing their contents into the extracellular space as exosomes. In this study, we analyzed the physical characteristics of EVs from 200 nm to 800 nm in diameter which we refer to as m/lEVs as per the MISEV2018 guidelines [9].

Recently, the clinical relevance of EVs has attracted significant attention. In particular, m/lEVs play an important role in tumor invasion [10]. m/lEVs in blood act as a coagulant factor and have been associated with sickle cell disease, sepsis, thrombotic thrombocytopenic purpura, and other diseases [11]. A possible role for urinary m/lEVs in diabetic nephropathy was also reported [12]. In recent years, the clinical applications of exosomes have advanced, and methods for detecting exosomes in patients with colorectal cancer have been developed [13]. However, because characterization of exosomes is analytically challenging, determining the cells and tissues from which exosomes are derived can be difficult. m/lEVs are generated differently from exosomes [14] but are similar in size and contain many of the same surface antigens. It is widely hypothesized that complete separation of exosomes and m/lEVs is likely to be a major challenge, and more effective techniques to purify and characterize m/lEVs would be extremely valuable. In this study, we focused on m/lEVs in plasma and urine, which are representative body fluids in clinical laboratories. We first developed a purification method for m/lEVs based on differentiation centrifugation. Second, we characterized m/lEVs by flow cytometry and mass spectrometry analysis and described the basic properties of m/lEV subpopulations in blood and urine.

## Materials and Methods

### Antibodies and other reagents

The following monoclonal antibodies against human surface antigens were used in this study: anti-CD5-Brilliant Violet 510 (clone: L17F12), anti-CD15-PerCP (clone: W6D3), anti-CD41-APC/Cy7 (clone: HIP8), anti-CD45-PE/Cy7 (clone: HI30), anti-CD59-PE (clone: p282), anti-CD61-PerCP (clone: VI-PL2), anti-CD105-Brilliant Violet 421 (clone: 43A3), anti-CD146-PE (clone: P1H12), anti-CD235a-APC/Cy7 (clone: HI264), anti-CD10-APC/Cy7 (clone: HI10a), anti-CD13-Brilliant Violet 421 (clone: WM15), anti-CD26-PE (clone: BA5b), anti-CD227 (MUC1)- PE/Cy7 (clone: 16A). All antibodies were purchased from BioLegend (San Diego, CA). FITC-conjugated Annexin V was purchased from BD Biosciences (New Jersey, USA). We used the SPHERO™ Nano Fluorescent Particle Size Standard Kit, Yellow (diameters 0.22, 0.45, 0.88 and 1.35 μm) from Spherotech Inc. for size validation. Normal mouse IgG was purchased from Wako Chemicals (Tokyo, Japan). APC-conjugated normal mouse IgG was produced using the Mix-n-Stain™ APC Antibody Labeling kit from Biotium Inc. Dithiothreitol (DTT) was purchased from Wako Chemicals (Tokyo, Japan). We conducted phase transfer surfactant experiments using “MPEX PTS Reagents for MS” purchased from GL Sciences Inc. and “Trypsin, TPCK Treated” purchased from Thermo Fisher Scientific. Iodoacetamide was purchased from Wako Chemicals (Tokyo, Japan).

### Samples

All studies were approved by the Institutional Review Board of the Kyushu University Hospital, Kyushu University (29-340). Blood samples were collected from 20 male and female participants (23-48 years of age) who were apparently healthy. Samples were collected using a 22-gauge butterfly needle and a slow-fill syringe. After discarding the initial 2-3 mL, blood was dispensed into collection tubes containing ethylenediamine tetra acetic acid (EDTA) (1.6 mg/mL blood). Urine was collected from 20 male healthy subjects (23-46 years of age). The first morning void urine was used for the experiments. The urine samples were collected in a sterile container. These experiments followed the International Society for EVs (ISEV) proposed Minimal Information for Studies of Extracellular Vesicles (“MISEV”) guidelines for the field in 2018 [9].

### Isolation of plasma m/lEVs

Essentially platelet-free plasma (PFP) was prepared from EDTA-treated blood by double centrifugation at 2,330 ×*g* for 10 min. To assess residual platelets remaining in this sample, we measured platelet number using the ADVIA 2120i Hematology System (SIEMENS Healthineers, Erlangen Germany). The number of platelets in this sample was below the limit of detection (1×10^3^ cells/μL). We used a centrifugation method to obtain m/lEVs. In an effort to ensure our approach could be applied to clinical testing, we chose a simple and easy method for pretreatment. In an ISEV position paper [15], Thery’s group referred to vesicles sedimenting at 100,000 ×*g* as “small EVs” rather than exosomes, those pelleting at intermediate speed (lower than 20,000 ×*g*) as “medium EVs” (including microvesicles and ectosomes) and those pelleting at low speed (e.g., 2000 ×*g*) as “large EVs”. Because these definitions are less biologically meaningful but more experimentally tractable than previously-applied exosome/microvesicle definitions, we attempted biological characterization through subsequent shotgun and flow cytometry analysis.

In flow cytometric analysis, the volume of PFP used in each assay was 0.6 mL from each donor. In nanoparticle tracking analysis (NTA) and electron microscopy, the volume of PFP used was 3 mL. Samples were independent and were treated individually prior to each measurement. PFP was centrifuged at 18,900 ×*g* for 30 min in a fixed-angle rotor. The m/lEV pellet obtained after centrifugation was reconstituted by vortex mixing (1-2 min) with an equivalent volume of Dulbecco’s phosphate-buffered saline (DPBS), pH 7.4. The solution was centrifuged at 18,900 ×*g* for 30 min again and the supernatant was discarded.

### Isolation of urinary m/lEVs

For isolation of urinary m/lEVs, we modified a urinary exosome extraction protocol [16]. The centrifugation conditions were identical for plasma and urine so that the size and the density of m/lEVs were similar, enabling comparison of plasma and urinary m/lEVs.

In flow cytometric analysis, the volume of urine used for each assay was 1.2 mL from each donor. In NTA and electron microscopy, the volume of urine used was 15 mL. Samples were independent and were treated individually prior to each measurement. Collected urine was centrifuged at 2,330 ×*g* for 10 min twice. The supernatant was centrifuged at 18,900 ×*g* for 30 min in a fixed-angle rotor. The m/lEV pellet obtained from centrifugation was reconstituted by vortex mixing (1-2 min) with 0.2 mL of DPBS followed by incubation with DTT (final concentration 10 mg/mL) at 37°C for 10-15 min. The samples were centrifuged again at 18,900 ×*g* for 30 min and the supernatant was discarded. Addition of DTT, a reducing agent, reduced the formation of Tamm-Horsfall protein (THP) polymers. THP monomers were removed from m/lEVs after centrifugation. DTT-containing DPBS solutions were filtered through 0.1-μm filters (Millipore).

### Flow cytometric analysis of m/lEVs

After resuspending m/lEV pellets in 60 μL of DPBS, we added saturating concentrations of several labelled antibodies, Annexin V and normal mouse IgG and incubated the tubes in the dark, without stirring, for 15-30 min at room temperature. In one case, we added labelled antibodies directly to 60 μL of PFP for staining (Supplementary Fig. S1). We resuspended stained fractions in Annexin V binding buffer (BD Biosciences: 10 mM HEPES, 0.14 mM NaCl, 2.5 mM CaCl_2_, pH 7.4) for analysis by flow cytometry. DPBS and Annexin V binding buffer were filtered through 0.1-μm filters (Millipore). Flow cytometry was performed using a FACSVerse™ flow cytometer (BD Biosciences). The flow cytometer was equipped with 405 nm, 488 nm and 638 nm lasers to detect up to 13 fluorescent parameters. The flow rate was 12 μL/min. Forward scatter voltage was set to 381, side scatter voltage was set to 340, and each threshold was set to 200. Excitation (Ex.) and emission (Em.) wavelengths as well as voltages used for each fluorophore were as follows: FITC, Ex. 488 nm, Em. 527/32 nm, voltage 442; PE, Ex. 488 nm, Em. 586/42 nm, voltage 411; PerCP, Ex. 488 nm, Em. 700/54 nm, voltage 556; PE/Cy7, Ex. 488 nm, Em. 783/56 nm, voltage 564.3, APC: Ex. 640 nm, Em. 660/10 nm, voltage 538.2; APC/Cy7, Ex. 640 nm, Em. 783/56 nm, voltage 584.8; Brilliant Violet 421, Ex. 405 nm, Em. 448/45 nm, voltage 538.2; and Brilliant Violet 510, Ex. 405 nm, Em. 528/45 nm, voltage 540. Flow cytometry was performed using FACSuite™ software (BD Biosciences) and data were analyzed using FlowJo software.

### Nanoparticle tracking analysis (NTA)

NTA measurements were performed using a NanoSight LM10 (NanoSight, Amesbury, United Kingdom). After resuspending mEV pellets in 50 μL of DPBS, samples were diluted eight-fold (plasma) and 100-fold (urinary) with PBS prior to measurement. Particles in the laser beam undergo Brownian motion and videos of these particle movements are recorded. NTA 2.3 software then analyses the video and determines the particle concentration and the size distribution of the particles. Twenty-five frames per second were recorded for each sample at appropriate dilutions with a “frames processed” setting of 1500. The detection threshold was set at “7 Multi” and at least 1,000 tracks were analyzed for each video.

### Electron microscopy

For immobilization, we added 100 μL of PBS and another 100 μL of immobilization solution (4% paraformaldehyde, 4% glutaraldehyde, 0.1 M phosphate buffer, pH 7.4) to m/lEV pellets. After stirring, we incubated at 4°C for 1 h. For negative staining, the samples were adsorbed to formvar film-coated copper grids (400 mesh) and stained with 2% phosphotungstic acid, pH 7.0, for 30 s. For observation and imaging, the grids were observed using a transmission electron microscope (JEM-1400Plus; JEOL Ltd., Tokyo, Japan) at an acceleration voltage of 100 kV. Digital images (3296 × 2472 pixels) were taken with a CCD camera (EM-14830RUBY2; JEOL Ltd., Tokyo, Japan).

### Protein digestion

We used approximately 50 mL of pooled healthy plasma and 100 mL of pooled healthy male urine from five healthy subjects for digestion of m/lEVs. PFP was prepared from EDTA-treated blood by double centrifugation at 2,330 ×*g* for 10 min. We pooled 50 mL of PFP and divided this sample into 10-mL aliquots in conical tubes. The PFP in each tube was centrifuged at 18,900 ×*g* for 30 min in a fixed-angle rotor. The m/lEV pellet obtained after centrifugation was reconstituted by vortex mixing (1-2 min) with 10 mL of DPBS. The solution was centrifuged again at 18,900 ×g for 30 min and the supernatant was discarded. We repeated these washing steps three times to reduce levels of contaminating free plasma proteins and small EVs for shotgun analysis. After the last centrifugation, we removed supernatants and froze the samples.

Collected urine was centrifuged at 2,330 ×*g* for 10 min twice. We pooled the supernatants from 50-mL samples and divided them into 10-mL conical tubes. The supernatant in each tube was centrifuged at 18,900 ×*g* for 30 min in a fixed-angle rotor. The m/lEV pellet obtained from centrifugation was reconstituted by vortex mixing (1-2 min) with 15 mL of DPBS followed by incubation with DTT (final concentration 10 mg/mL) at 37°C for 10-15 min. The solution was centrifuged again at 18,900 ×*g* for 30 min and the supernatant including DTT was discarded. The m/lEVs pellet obtained after centrifugation was reconstituted by vortex mixing (1-2 min) with 10 mL of DPBS. The solution was centrifuged at 18,900 ×*g* for 30 min again and the supernatant was discarded. We repeated these washing steps twice to reduce levels of contaminating free urinary proteins and small EVs for shotgun analysis. We removed supernatants and froze the samples.

Samples were digested using a phase transfer surfactant-aided procedure [17]. The precipitated frozen fractions of plasma and urine were thawed at 37°C, and then m/lEVs were solubilized in 250 μL of lysis buffer containing 12 mM sodium deoxycholate and 12 mM sodium lauroyl sarcosinate in 100 mM Tris·HCl, pH 8.5. After incubating for 5 min at 95°C, the solution was sonicated using an ultrasonic homogenizer. Protein concentrations of the solutions were measured using a bicinchoninic acid assay (Pierce™ BCA Protein Assay Kit; Thermo Fisher Scientific).

Twenty microliters of the dissolved pellet (30 μg protein) were used for protein digestion. Proteins were reduced and alkylated with 1 mM DTT and 5.5 mM iodoacetamide at 25°C for 60 min. Trypsin was added to a final enzyme:protein ratio of 1:100 (wt/wt) for overnight digestion. Digested peptides were acidified with 0.5% trifluoroacetic acid (final concentration) and 100 μL of ethyl acetate was added for each 100 μL of digested m/lEVs. The mixture was shaken for 2 min and then centrifuged at 15,600 ×*g* for 2 min to obtain aqueous and organic phases. The aqueous phase was collected and desalted using a GL-Tip SDB column (GL Sciences Inc).

### LC-MS/MS analysis

Digested peptides were dissolved in 40 μL of 0.1% formic acid containing 2% (v/v) acetonitrile and 2 μL were injected into an Easy-nLC 1000 system (Thermo Fisher Scientific). Peptides were separated on an Acclaim PepMap™ RSLC column (15 cm × 50 μm inner diameter) containing C18 resin (2 μm, 100 Å; Thermo Fisher Scientific), and an Acclaim PepMap™ 100 trap column (2 cm× 75 μm inner diameter) containing C18 resin (3 μm, 100 Å; Thermo Fisher Scientific). The mobile phase consisted of 0.1% formic acid in ultrapure water (buffer A). The elution buffer was 0.1 % formic acid in acetonitrile (buffer B); a linear 200 min gradient from 0%-40% buffer B was used at a flow rate of 200 nL/min. The Easy-nLC 1000 was coupled via a nanospray Flex ion source (Thermo Fisher Scientific) to a Q Exactive Orbitrap (Thermo Fisher Scientific). The mass spectrometer was operated in data-dependent mode, in which a full-scan MS (from 350 to 1,400 m/z with a resolution of 70,000, automatic gain control (AGC) 3E+06, maximum injection time 50 ms) was followed by MS/MS on the 20 most intense ions (AGC 1E+05, maximum injection time 100 ms, 4.0 m/z isolation window, fixed first mass 100 m/z, normalized collision energy 32 eV).

### Proteome Data Analysis

Raw MS files were analyzed using Proteome Discoverer software version 1.4 (Thermo Fisher Scientific) and peptide lists were searched against the Uniprot Proteomes-Homo sapiens FASTA (Last modified November 17, 2018) using the Sequest HT algorithm. Initial precursor mass tolerance was set at 10 ppm and fragment mass tolerance was set at 0.6 Da. Search criteria included static carbamidomethylation of cysteine (+57.0214 Da), dynamic oxidation of methionine (+15.995 Da) and dynamic acetylation (+43.006 Da) of lysine and arginine residues.

### Gene ontology analysis and gene enrichment analysis

We conducted GO analysis using DAVID (https://david.ncifcrf.gov) to categorize the proteins identified by shotgun analysis. We uploaded the UNIPROT_ACCESSION No. for each protein and showed the categorization of cellular components from the results of “Functional Annotation Chart.”

We used Metascape (http://metascape.org/gp/index.html#/main/step1) for gene enrichment analysis. We uploaded the UNIPROT_ACCESSION No. for each protein, selected human as species and performed an express analysis. For each gene list, pathway and process enrichment analysis was carried out using the following ontology sources: KEGG Pathway, GO Biological Processes, Reactome Gene Sets, Canonical Pathways and CORUM. The entire human genome was used as the enrichment background.

### Dipeptidyl peptidase IV (DPP4) activity assay

DPP4 activity was measured in the plasma and urine of six individuals (different from plasma donors). The method was previously published in part [18]. DPP4 activity was measured via the fluorescence intensity of 7-amino-4-methylcoumarin (AMC) after its dissociation from the synthetic substrate (Gly-Pro-AMC • HBr) catalyzed by DPP4. Experiments were performed in 96-well black plates. Titrated AMC was added to each well to prepare a standard curve. Fluorescence intensity was measured after incubating substrate with urine samples for 10 min. The enzyme reaction was terminated by addition of acetic acid. The fluorescence intensity (Ex. = 380 nm and Em. = 460 nm) was measured using Varioskan Flash (Thermo Fisher Scientific). DPP4 activity assays were performed by Kyushu Pro Search LLP (Fukuoka, Japan).

## Results

### Isolation and characterization of m/lEVs from plasma and urine

The workflow for the isolation and enrichment of m/lEVs for proteomic and flow cytometric analyses is illustrated in Fig.1A and 1B. m/lEVs from human plasma samples were isolated by high-speed centrifugation, an approach used in previous studies [19]. For isolation of m/lEVs from urine, DTT, a reducing agent, was used to remove THP polymers because these non-specifically interact with IgGs.

Transmission electron microscopy revealed that almost all m/lEVs were small, closed vesicles with a size of approximately 200 nm that were surrounded by lipid bilayer (Fig. 1C-H). In plasma, we observed EVs whose membranes were not stained either inside or on the surface (Fig. 1C, 1D); we also observed EVs whose forms were slightly distorted (Fig. 1E). In urine, a group of EVs with uneven morphology and EVs with interior structures were observed (Fig. 1F-1H). Apoptotic bodies, cellular debris, and protein aggregates were not detected.

Next, we measured EV particle size by NTA. NTA of plasma and urine m/lEVs revealed a peak at ~200 nm containing small EVs (S1 Fig.). No EVs with diameters greater than 800 nm were observed by NTA and flow cytometry can detect only EVs with diameters larger than 200 nm. Together, these data suggested that we characterized m/lEVs between 200 nm and 800 nm in diameter from plasma and urine by flow cytometry analysis.

Side-scatter events from size calibration beads of (diameters: 0.22, 0.45, 0.88 and 1.35 μm) were resolved from instrument noise using a FACS Verse flow cytometer (S2 Fig. A). Inspection of the side-scatter plot indicated that 0.22 μm was the lower limit for bead detection. More than 90% of m/lEVs isolated from plasma and urine showed side-scatter intensities lower than those of 0.88-μm calibration beads (Fig. 2A-D). m/lEVs were heterogeneous in size, with diameters ranging from 200-800 nm in plasma and urine (Fig. 2A-D). Fluorescently-labeled mouse IgG was used to exclude nonspecific IgG-binding fractions (S2 Fig. B and C). In this experiment, we characterized m/lEVs with diameters ranging from 200-800 nm. Using these methods, we observed an average of 8×10^5^m/lEVs in each mL of plasma.

**Fig 1.**
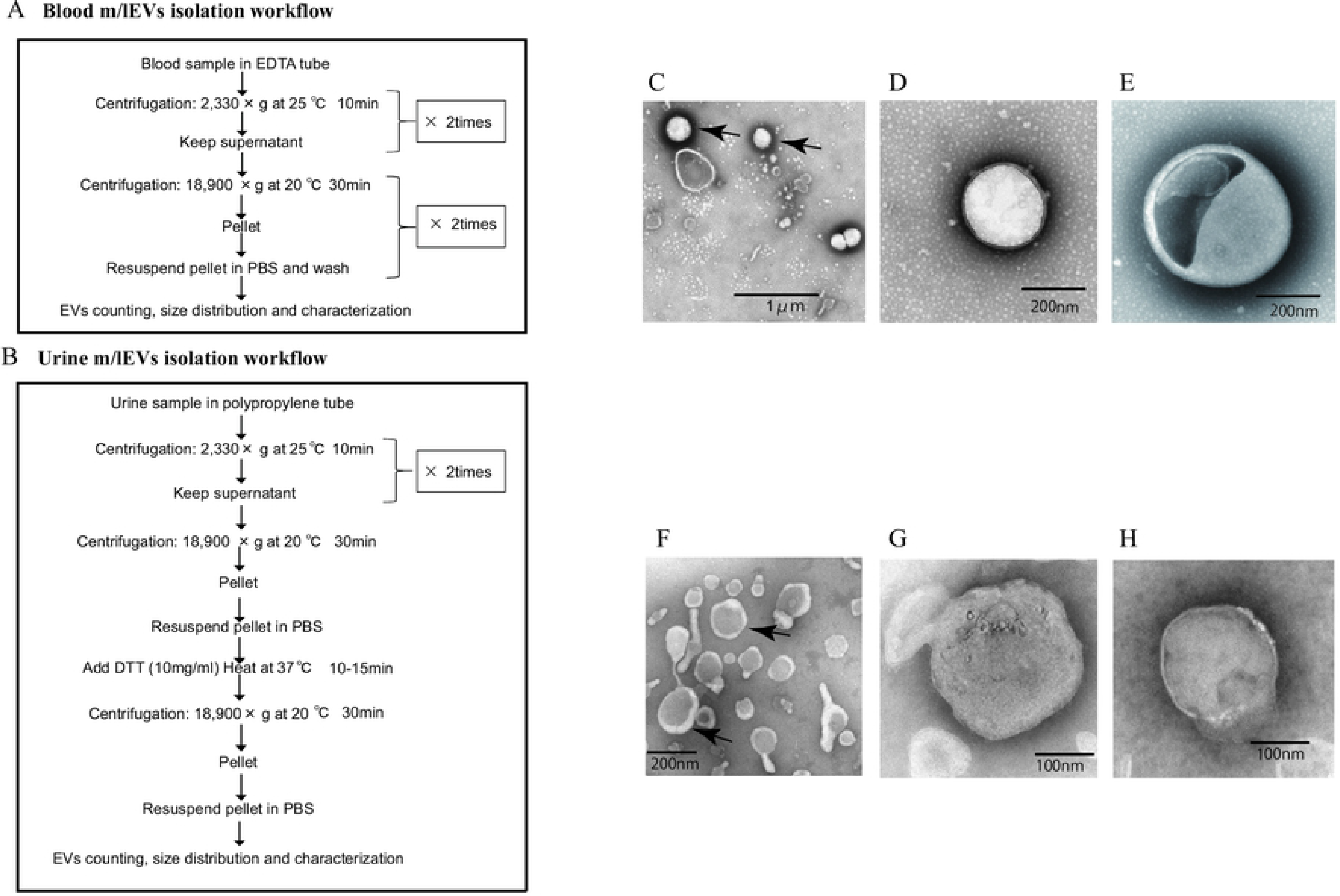
Isolation of m/lEVs from plasma and urine using differential centrifugation. (A and B) Workflow of plasma (A) and urine (B) m/lEV isolation and sample preparation for flow cytometry analysis. (C-H) Isolated m/lEVs from plasma (C-E) and urine (F-H) were visualized by transmission electron microscopy. Arrow indicates representative m/lEVs (C and F). Microscopy was used to identify EV-like particles based on the size (100-400 nm) and shape (round) of the vesicles. The scale bar is shown.

**Fig 2.**
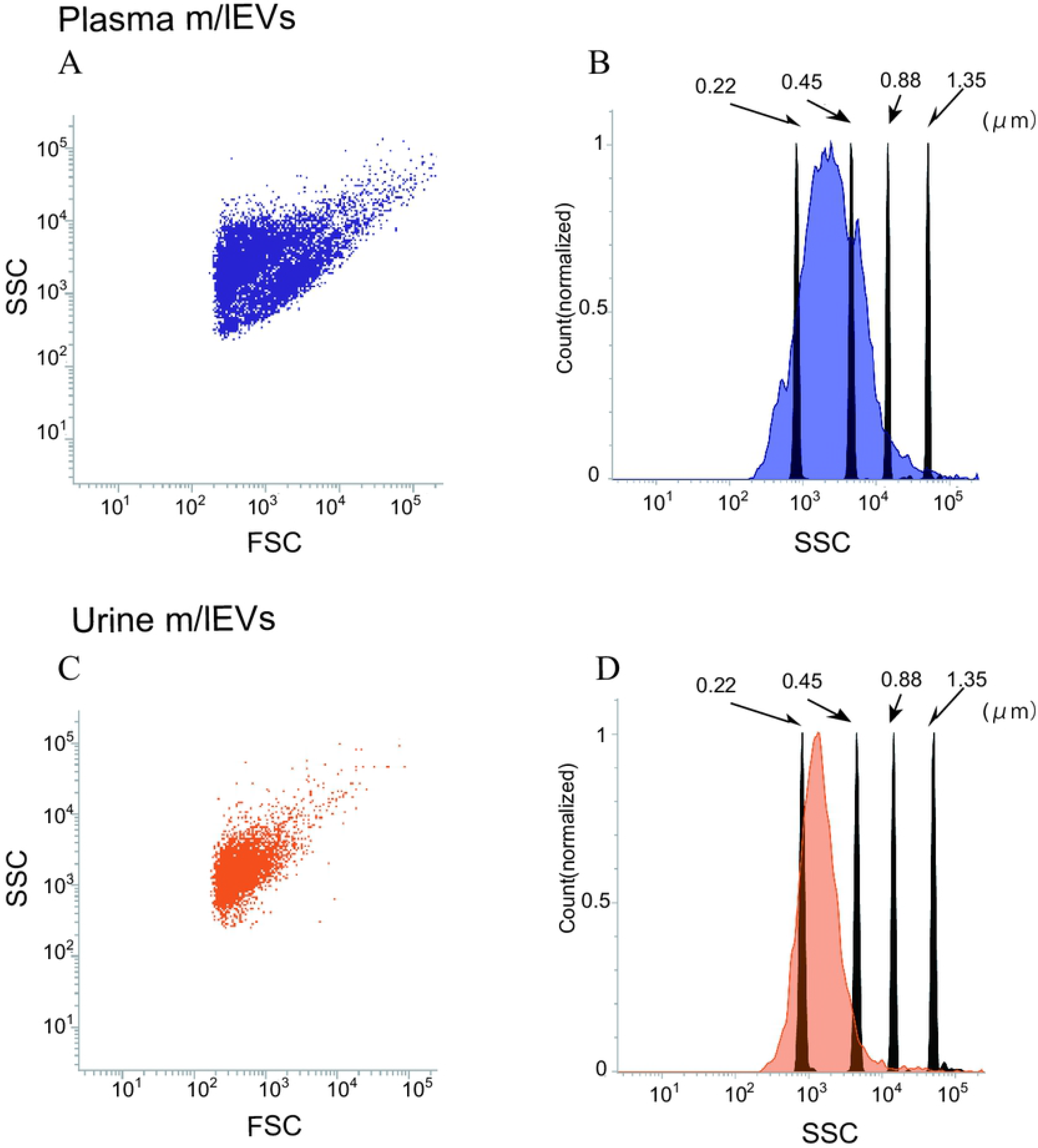
Flow cytometric analysis of plasma and urine m/lEVs. (A and B) Analysis of plasma m/lEVs by flow cytometry. Forward and side scatter (SSC) were measured for plasma m/lEVs (A). The SSC distribution of plasma m/lEVs is shown as a histogram (indigo blue) compared with standard polystyrene beads (black histogram) (B). (C and D) Analysis of urine m/lEVs by flow cytometry. Forward and side scatter (SSC) were measured for urine m/lEVs (C). The SSC distribution of urinary m/lEVs is shown as a histogram (orange) compared with standard polystyrene beads (black histogram) (D).

### Shotgun proteomic analysis of plasma and urine EVs

To analyze the protein components of m/lEVs present in plasma and urine of five healthy individuals, we performed LC-MS/MS proteomic analysis. A total of 621 and 1,793 proteins were identified in m/lEVs from plasma and urine, respectively (Fig. 3A and S1 Table and S2 Table). Scoring counts using the SequestHT algorithm for the top 20 most abundant proteins are shown in Tables 1 and 2. We detected cytoskeleton-related protein such as actin, ficolin-3 and filamin and cell-surface antigen such as CD5, band3 and CD41 in plasma. We also identified actin filament-related proteins such as ezrin, radixin, ankylin and moesin which play key roles in cell surface adhesion, migration and organization. In urine, several types of peptidases (membrane alanine aminopeptidase or CD13; neprilysin or CD10; DPP4 or CD26) and MUC1 (mucin 1 or CD227) were detected in high abundance, and these proteins were used to characterize EVs by flow cytometric analysis (Table 2 and S2 Table). We demonstrated that the isolated m/lEVs showed high expression of tubulin and actinin, while the tetraspanins CD9 and CD81 that are often used as exosome markers were only weakly identified. These data suggest that m/lEVs differ from small EVs including exosomes (S3 Table).

As shown in Fig. 3A and S3 Fig., about 10% of urinary EVs proteins were also identified in plasma EVs. Urinary EVs in the presence of blood contaminants were also observed in previous studies [20]. These result suggest that m/lEVs in plasma were excreted in the urine via renal filtration and not reabsorbed. Gene ontology analysis of the identified proteins indicated overall similar cellular components in plasma and urine m/lEVs (Fig. 3B). The results of gene set enrichment analysis by metascape are shown for plasma and urine m/lEVs (Fig. 3C, D and S4 Table and S5 Table). The most commonly-observed functions in both plasma and urine were “regulated exocytosis”, “hemostasis” and “vesicle-mediated transport”. In plasma, several functions of blood cells were observed, including “complement and coagulation cascades” and “immune response”. Moreover, analysis of urinary EVs showed several characteristic functions including “transport of small molecules”, “metabolic process” and “cell projection assembly”. This may reflect the nature of the kidney, the urinary system and tubular villi. Plasma-derived m/lEVs showed no evidence of exosome-related proteins such as TSG101, Alix and CD63, suggesting that our differential centrifugation method can efficiently yield m/lEVs (S3 Table). These data demonstrate the power of data-driven biological analyses.

**Fig 3.**
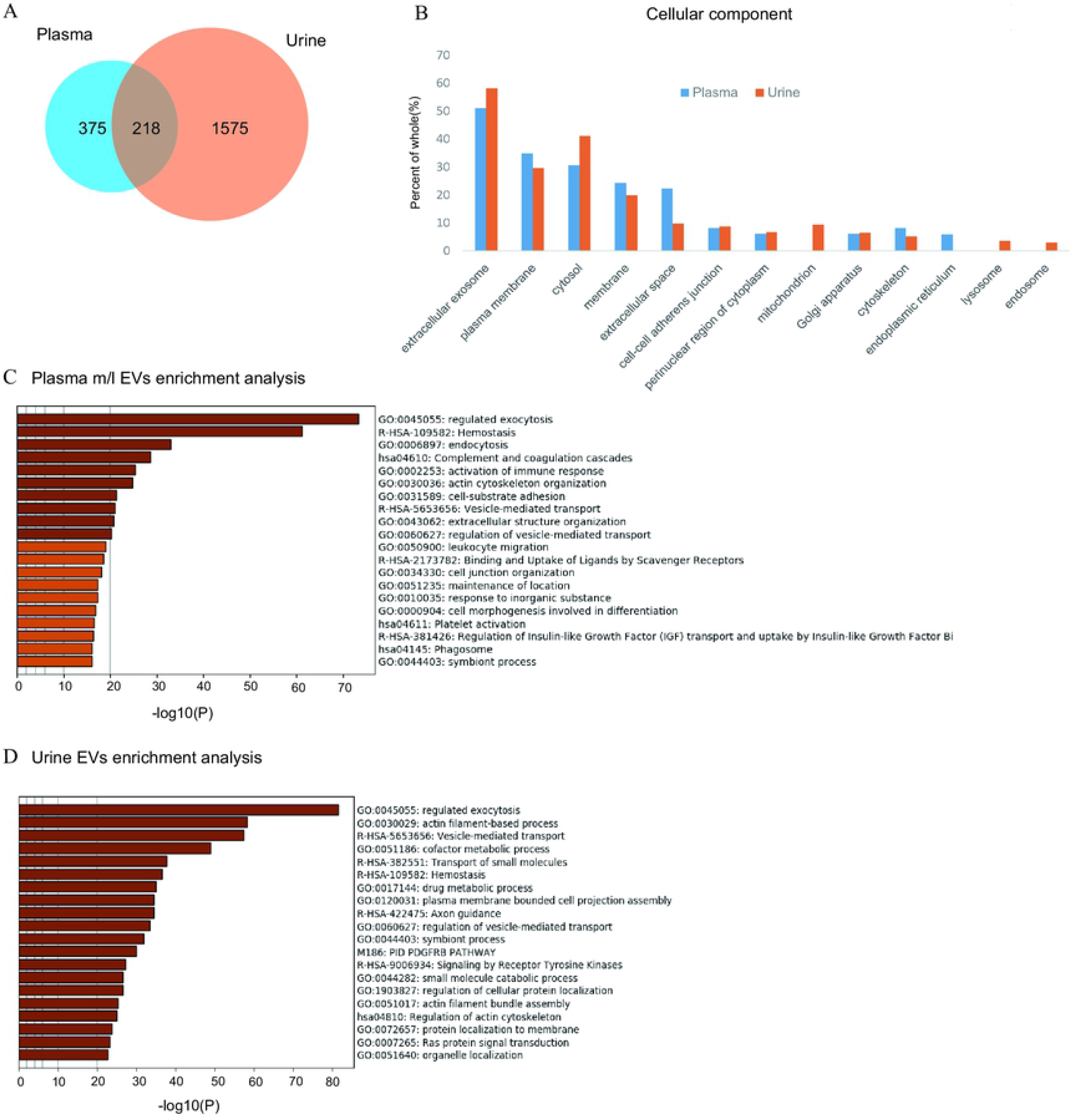
Shotgun proteomic analysis of plasma and urine m/lEVs. A. Protein extracts of m/lEVs isolated from plasma and urine were analyzed by LC-MS/MS. A total of 593 and 1793 proteins from plasma and urine, respectively, were detected. Detailed lists of proteins are shown in S1 Table and S2 Table. B. GO (gene ontology) cellular components are shown for m/lEVs isolated from plasma and urine using the DAVID program. Among the detected proteins, the gene list used for DAVID analysis included 588 proteins (plasma) and 1786 proteins (urine). The vertical axis shows the percentage of proteins from the full gene list categorized into each GO term. For example, for extracellular exosomes (plasma), the categorized count was 301 of 588 proteins. (C and D) Top 20 clusters from the Metascape pathway (http://metascape.org/) enrichment analysis for m/lEVs in plasma (C) and urine (D). Lengths of bars represent log10 (P values) based on the best-scoring term within each cluster. Among all detected proteins, 535 (plasma) and 1767 (urine) genes were recognized as unique for enrichment analysis. For each gene list, pathway and process enrichment analysis was carried out using the following ontology sources: KEGG Pathway, GO Biological Processes, Reactome Gene Sets, Canonical Pathways and CORUM.

**Table 1.** Twenty most abundant proteins identified in plasma m/lEVs. Proteins in bold text indicate antigens identified using flow cytometry. This table excluded immunoglobulin-related proteins and albumin.

| Protein name | Uniprot AC | Score | Sequence coverage | #PSMs |
| --- | --- | --- | --- | --- |
| Ficolin-3 | O75636 | 8772 | 68 | 3428 |
| Hemoglobin subunit alpha | P69905 | 1735 | 70 | 725 |
| Actin, cytoplasmic 1 | P60709 | 927 | 69 | 333 |
| Hemoglobin subunit beta | P68871 | 602 | 79 | 235 |
| Actin, gamma-enteric smooth muscle | P63267 | 500 | 32 | 210 |
| Talin-1 | Q9Y490 | 368 | 50 | 173 |
| Filamin-A | P21333 | 353 | 40 | 133 |
| Spectrin alpha chain, erythrocytic 1 | P02549 | 321 | 42 | 126 |
| Myosin-9 | P35579 | 317 | 38 | 110 |
| Mannan-binding lectin serine protease 1 | P48740 | 308 | 35 | 103 |
| Band 3 anion transport protein | P02730 | 277 | 38 | 137 |
| Beta-actin-like protein 2 | Q562R1 | 252 | 24 | 101 |
| Hemoglobin subunit delta | P02042 | 215 | 64 | 80 |
| Spectrin beta chain, erythrocytic | P11277 | 183 | 28 | 88 |
| Complement C1q subcomponent subunit C | P02747 | 170 | 24 | 49 |
| Ankyrin-1 | P16157 | 164 | 28 | 63 |
| **CD5 antigen-like** | O43866 | 164 | 52 | 60 |
| **Integrin alpha-IIb (CD41)** | P08514 | 129 | 25 | 44 |
| Complement C1q subcomponent subunit B | P02746 | 122 | 41 | 56 |
| Deaminated glutathione amidase | Q86X76 | 118 | 6 | 44 |

**Table 2.** Twenty most abundant proteins identified in urinary m/lEVs. Proteins in bold text indicate antigens identified using flow cytometry

| Protein name | Uniprot AC | Score | Sequence coverage | #PSMs |
| --- | --- | --- | --- | --- |
| Actin, cytoplasmic 1 | P60709 | 1618 | 67 | 566 |
| **Neprilysin** | P08473 | 1194 | 50 | 495 |
| Uromodulin | P07911 | 821 | 44 | 323 |
| Solute carrier family 12 member 1 | Q13621 | 720 | 32 | 251 |
| Alpha-enolase | P06733 | 708 | 79 | 339 |
| Moesin | P26038 | 548 | 73 | 208 |
| Ezrin | P15311 | 544 | 56 | 213 |
| **Aminopeptidase N** | P15144 | 486 | 43 | 214 |
| Actin, gamma-enteric smooth muscle | P63267 | 476 | 28 | 197 |
| Pyruvate kinase PKM | P14618 | 447 | 64 | 130 |
| Voltage-dependent anion-selective channel protein 1 | P21796 | 446 | 74 | 159 |
| Radixin | P35241 | 364 | 56 | 144 |
| Tyrosine-protein phosphatase non-receptor type 13 | Q12923 | 363 | 38 | 150 |
| Triosephosphate isomerase | P60174 | 325 | 80 | 108 |
| Multidrug resistance protein 1 | P08183 | 324 | 38 | 132 |
| Guanine nucleotide-binding protein G(I)/G(S)/G(T) subunit beta-2 | P62879 | 316 | 50 | 107 |
| Guanine nucleotide-binding protein G(I)/G(S)/G(T) subunit beta-1 | P62873 | 306 | 53 | 112 |
| V-type proton ATPase catalytic subunit A | P38606 | 298 | 57 | 137 |
| Superoxide dismutase [Cu-Zn] | P00441 | 285 | 91 | 63 |
| Epidermal growth factor receptor kinase substrate 8-like protein 2 | Q9H6S3 | 277 | 43 | 92 |

### Characterization of plasma EVs by flow cytometry

Next, we characterized m/lEVs in plasma by flow cytometry using antibodies against several surface antigens and Annexin V. We first excluded dart-out and agglomerating m/lEVs using the pulse width of the side-scatter plot. Furthermore, to eliminate nonspecific adsorption, we excluded the mouse IgG-positive fraction. (S4 Fig.) We characterized positive m/lEVs using labeled antibodies against nine surface antigens and Annexin V (Fig. 4A-C). Among these nine antigens, six [CD5, CD41 (integrin alpha-IIb), CD45 (receptor-type tyrosine-protein phosphatase C), CD61 (integrin β-3) and CD59, and CD235a (glycophorin A)] were also detected by shotgun proteomic analysis. We did not detect the exosomal marker proteins TSG101 or Alix in m/lEVs by flow cytometric analysis (data not shown).

To characterize m/lEVs derived from erythrocytes, T and B cells, macrophages/monocytes, granulocytes, platelets and endothelial cells, we selected nine antigens described in Fig. 4A. Two or more antigens were used for characterization of m/lEVs: for example, CD59 and CD235a double-positive and CD45-negative m/lEVs were classified as erythrocyte-derived m/lEVs (S4 Fig.B). We also show Annexin V staining for the m/lEVs corresponding to these five classifications in Fig. 4B and C. We integrated these characterizations and assessed the distribution of EV classifications among ten healthy subjects (Fig. 4D). The results suggested that no major differences in the ratios of fractions in these ten subjects and thus these definitions may be used for pathological analysis.

We found that 10% and 35% of m/lEVs were derived from erythrocytes and platelets, respectively. However, only 0.5%, 0.6% and 0.1% of m/lEVs were derived from macrophages, leukocytes and endothelial cells, respectively suggesting that the ratio of m/lEVs of different cellular origins is dependent on the number of cells present in plasma (Fig. 4D). We also observed that most m/lEVs derived from erythrocytes, macrophages and vascular endothelium were Annexin V positive. By contrast, many Annexin V negative m/lEVs were identified among platelet- and T and B cell-derived m/lEVs (Fig. 4E). These results suggested that Annexin V positive m/lEVs are cell-type specific and that release mechanisms may differ among cell types.

**Fig 4.**
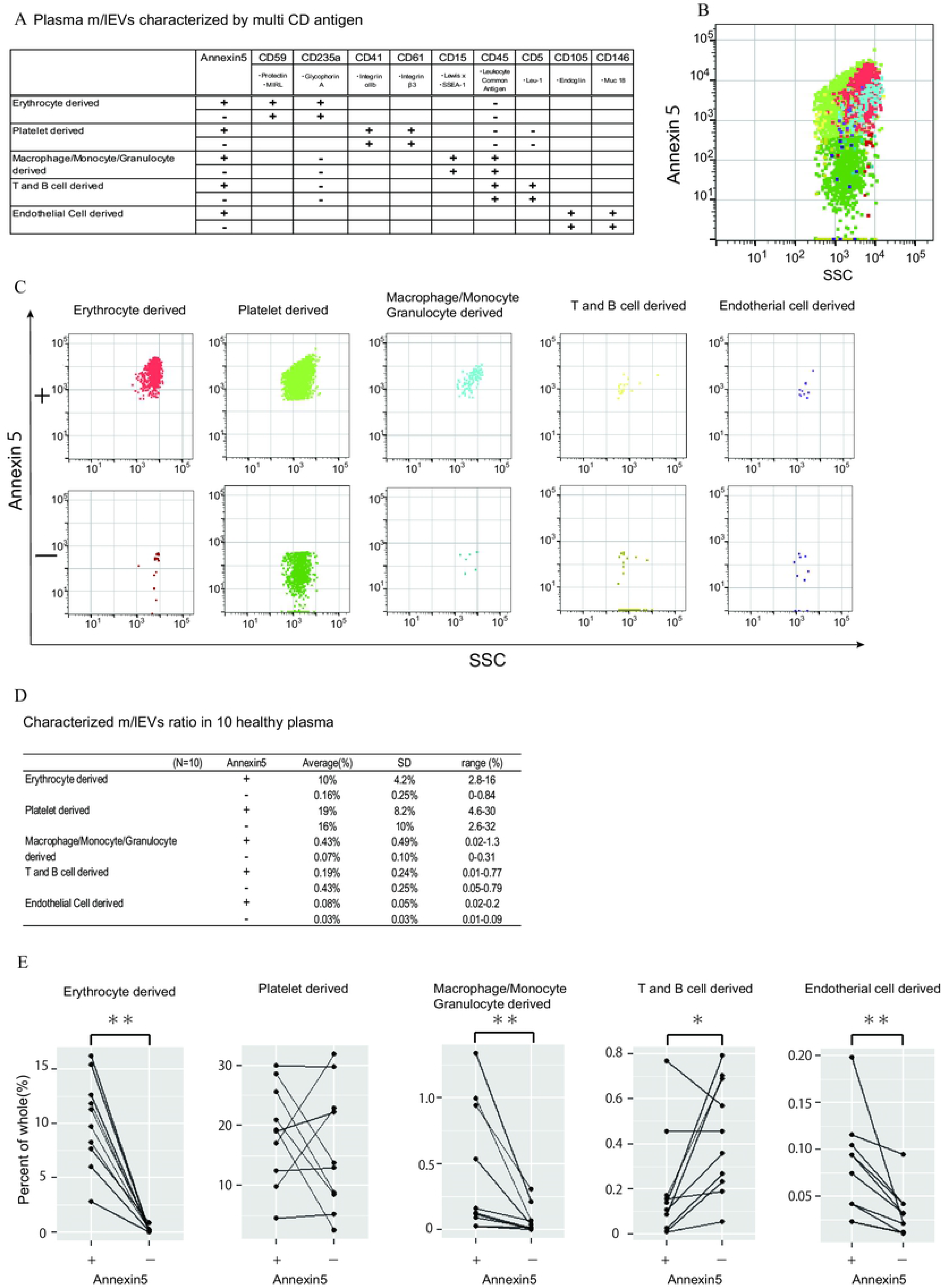
Characterization of plasma m/lEVs by flow cytometry. A. Two specific surface antigens were used to characterize the m/lEVs from each source by flow cytometry. m/lEVs were characterized by comparison with five types of blood cells using surface antigens and Annexin V staining. (B and C). Representative dot plots (SSC vs. Annexin V). Each plot was classified by staining for surface antigens and Annexin V (C) and an overall plot summarizing the data (B) is shown. D. Quantification of each m/lEV by flow cytometry analysis (n=10 for human healthy plasma, % of total events). E. Comparison of Annexin V staining for erythrocyte-, platelet-, macrophage-, T and B cell-, and endothelial cell-derived m/lEVs from ten healthy plasma samples. Comparisons were performed using the Wilcoxon signed-rank test (* p< 0.05, ** p < 0.01; a value of p < 0.05 was considered to indicate statistical significance).

### Characterization of urinary EVs by flow cytometry and enzyme activity assay

In urine, we first removed aggregated m/lEVs and residual THP polymers using labelled normal mouse IgG (S2 Fig. C). To characterize urinary m/lEVs, we used surface antigens detected by shotgun proteomic analysis including CD10 (neprilysin), CD13 (alanine aminopeptidase), CD26 (DPP4) and CD227 (MUC1) (Fig. 5A and B). Many m/lEVs in the observation area were triple-positive for CD10, CD13 and CD26, but negative for Annexin V (Fig. 5B, S5 Fig.). Furthermore, MUC1-positive EVs were both Annexin V positive and negative in roughly equivalent frequencies (Fig. 5B and Fig. 5D). These results suggested that m/lEVs containing peptidases were released by outward budding directly from the cilial membrane of renal proximal tubule epithelial cells. The results of integrating these characterizations and the distribution of EV classifications among ten healthy subjects are shown in Fig. 5C. These data indicated no major differences in the ratio among these populations, suggesting that our methodology was reliable for m/lEV analysis.

**Fig 5.**
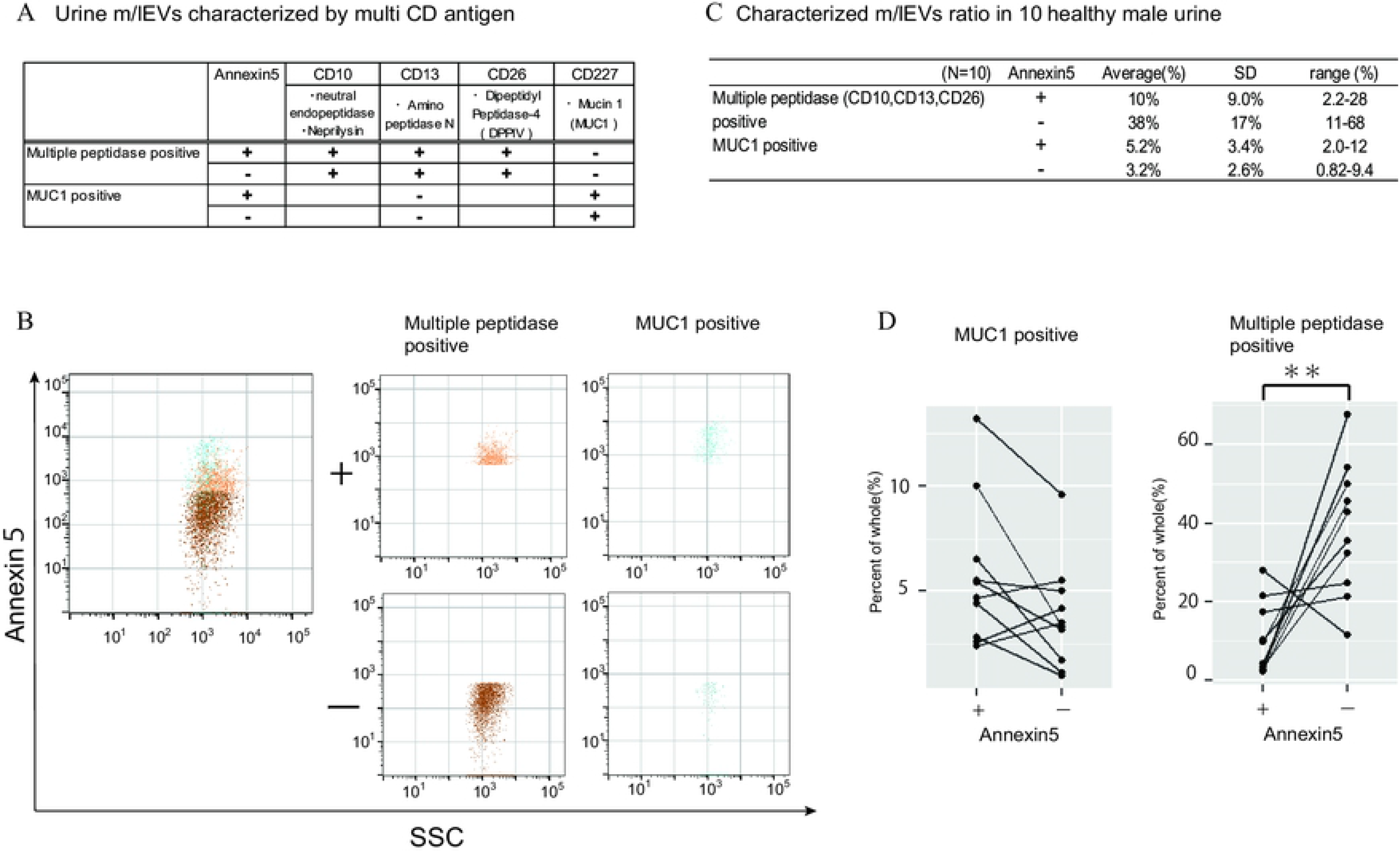
Characterization of urinary m/lEVs by flow cytometry. A. Characterization of urine m/lEVs was performed using surface antigens identified by the shotgun method. m/lEVs were characterized by comparison with two kinds of cells using surface antigen and Annexin V staining. B. Representative dot plots in the observation area (SSC vs Annexin V). Each plot was classified according to staining for multiple peptidases [CD10 (neprilysin), CD13 (alanine aminopeptidase), and CD26 (DPP4)] and CD277 (MUC-1). Events were further classified as Annexin V positive or negative. C. Quantification of m/lEVs by flow cytometry analyses (n=10 for male human healthy urine, % of total events). D. Comparison of Annexin V staining for MUC1-positive and multiple peptidase-positive m/lEVs in the urine of ten healthy male human samples. Comparisons were performed using the Wilcoxon signed-rank test (* P < 0.05, ** P < 0.01; a value of p < 0.05 was considered to indicate statistical significance).

**Fig 6.**
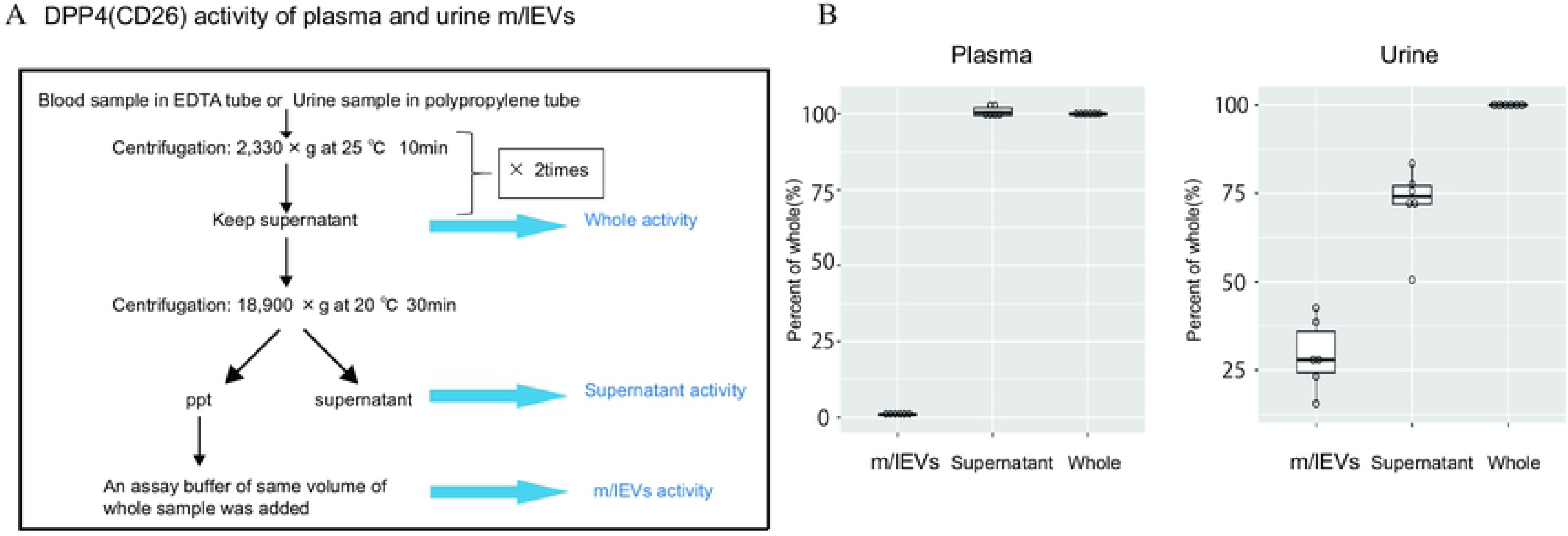
DPP4 enzymatic activity of plasma and urinary m/lEVs. A. DPP4 enzymatic activity was assessed for urinary m/lEVs. The workflow for fractionating m/lEVs by centrifugation is shown. B. Quantification of DPP4 activity in plasma and urine fractions (n=6 for human healthy plasma and male urine, % of whole). The percentage of enzyme activity measured for each fraction compared with whole activity is shown.

We next examined the peptidase enzyme activities of m/lEVs in plasma and urine from six individuals. We prepared three fractions: (i) “whole,” in which debris were removed after low speed centrifugation, (ii) “m/lEVs” and (iii) “free (supernatant)” both of which were obtained via high speed centrifugation (18,900 ×*g* for 30 min) (Fig. 6A). We found that more than 20% of DPP4 activity in whole urine was contributed by the EV fraction (Fig. 6B and S6 Fig.). By contrast, there was no peptidase activity associated with plasma m/lEVs (Fig. 6B). These results suggested that functional peptidase activity is present in m/lEVs in urine, which may be useful for pathological analysis.

## Discussion

In this study, we analyzed m/lEVs using various analytic techniques and found the following five major results. First, it was possible to characterize m/lEVs using multiple surface markers. Second, m/lEVs are likely to contain various cell-derived organelles. Third, m/lEVs bear functional enzymes with demonstrable enzyme activity on the vesicle surface. Fouth, there are differences in asymmetry of membrane lipids by derived cells. Finally, there was little variation m/lEVs in the plasma and urine of healthy individuals, indicating that our method is useful for identifying cell-derived m/lEVs in these body fluids.

We isolated m/lEVs from plasma and urine that were primarily 200-800 nm in diameter as shown by NTA and transmission electron microscopy. A large proportion of proteins detected in m/lEVs using shotgun proteomic analysis were categorized as plasma membrane proteins. Isolation of m/lEVs by centrifugation is a classical technique, but in the present study we further separated and classified the m/lEVs according to their cell types of origin by flow cytometry. The results indicated the validity of the differential centrifugation method [3, 21].

Pang et al. [22] reported that integrin outside-in signaling is an important mechanism for microvesicle formation, in which the procoagulant phospholipid phosphatidylserine (PS) is efficiently externalized to release PS-exposed microvesicles (MVs). These platelet-derived Annexin V positive MVs were induced by application of a pulling force via an integrin ligand such as shear stress. This exposure of PS allows binding of important coagulation factors, enhancing the catalytic efficiencies of coagulation enzymes. We observed that 50% of m/lEVs derived from leukocytes and platelets were Annexin V positive, suggesting that these cells are exposed to shear stress during blood flow and release PS-positive m/lEVs during activation, inflammation, and injury. It would be interesting to further investigate whether the ratio of Annexin V positive m/lEVs from platelets or leukocytes was an important diagnostic factor for inflammatory disease or tissue injury.

A previous report suggested that flow cytometry side-scatter values were suitable for observation of EVs less than 1000 nm in diameter, and in our study m/lEV size was measured by comparison with polystyrene beads [23, 24]. Based on the results of NTA analysis, small materials ≤100 nm in diameter are included in fractions isolated by centrifugation. In this study, we focused on m/lEVs more than 200 nm in diameter and characterized them by flow cytometry.

In urinary m/lEVs, we identified aminopeptidases such as CD10, CD13 and CD26 which are localized in proximal renal tubular epithelial cells. The functions of these proteins relating to exocytosis were categorized by gene enrichment analysis. The cilium in the kidney is the site at which a variety of membrane receptors, enzymes and signal transduction molecules critical to many cellular processes function. In recent years, ciliary ectosomes - bioactive vesicles released from the surface of the cilium - have attracted attention [25–27]. A good candidate mechanism for ciliary ectosome formation is membrane remodeling by ESCRT complexes, which include the charged multivesicular body proteins 2A, 2B, 4A, 4B, 4C, and 6 and vacuolar protein sorting proteins 4A and 4B [25, 28]. We also identified these proteins in proteomic analyses, suggesting that our isolation method was valid and the possibility that these proteins were biomarkers of kidney disease. Because triple peptidase positive m/lEVs were negative for Annexin V, the mechanism of budding from cells may not be dependent on scramblase. [25]

Platelet-derived m/lEVs are the most abundant population of extracellular vesicles in blood, and their presence [11] and connection with tumor formation were reported in a recent study [29]. In our study, platelet-derived EVs were observed in healthy subjects and had the highest abundance of Annexin V-positive EVs. In plasma, leukocyte-derived EVs were defined as CD11b/CD66b- or CD15-positive [30]. We characterized macrophage/monocyte/granulocyte- and T/B cell-derived EVs based on two specific CD antigens, and we confirmed that EVs derived from these cells were very rare. We also characterized small numbers of endothelial-derived EVs bearing CD146 and CD105 in healthy subjects. It was reported that endothelial-derived EVs were CD31-positive, Annexin V-positive and CD42b-negative, and that levels of these EVs were elevated in patients with systemic lupus erythematosus [31] and carotid artery disease [32]. Importantly, there was little variation in the cellular origins of m/lEVs in samples from ten healthy individuals, indicating that this method was useful for identifying cell-derived m/lEVs. We plan to examine m/lEVs differences in patients with these diseases in the near future. Erythrocyte-derived EVs were also characterized by their expression of CD235a and glycophorin A by flow cytometry [33, 34]. Because we observed that erythrocyte-derived EVs were elevated in hemolytic plasma (data not shown), any clinical applications should take into account their hemolytic presence.

We also characterized m/lEVs in urine. In kidneys and particularly in the renal tubule, CD10, CD13, CD26 can be detected in high abundance by immunohistochemical staining (website: The Human Protein Atlas). CD10/CD13-double positive labeling can be used for isolation and characterization of primary proximal tubular epithelial cells from human kidney [35]. The urinary exosome membrane protein, alanyl aminopeptidase or CD13, is a marker of infection in rats [36]. DPP4 (CD26) is a potential biomarker in urine for diabetic kidney disease and the presence of urinary m/lEV-bound DPP4 has been demonstrated [12]. The presence of peptidases on the m/lEV surface, and their major contribution to peptidase activity in whole urine [12], may suggest a functional contribution to reabsorption in the proximal tubules. These observations suggested that the ratio of DPP activity between m/lEVs and total urine can be an important factor in the diagnosis of kidney disease.

MUC1 can also be detected in kidney and urinary bladder by immunohistochemical staining (website: The Human Protein Atlas). Significant increases of MUC1 expression in cancerous tissue and in the intermediate zone compared with normal renal tissue distant from the tumor was observed [37]. High MUC1 expression was associated with a favorable prognosis in patients with bladder cancer [38]. In any case, MUC1-positive EVs are thought to be more likely to be derived from the tubular epithelium or the urothelium.

We made use of these findings to analyze the tissues and cells from which EVs were derived. Use of EVs as diagnostic reagents with superior disease and organ specificity for liquid biopsy samples is a possibility. This protocol will allow further study and in depth characterization of EV profiles in large patient groups for clinical applications. We are going to attempt to identify novel biomarkers by comparing healthy subjects and patients with various diseases.

## ACKNOWLEDGEMENTS

We thank lab members for reagents, discussions, and critical reading of the manuscript. We are grateful to LSI Medience Corporation for generous support during this study. This work was supported by a Grant-in-Aid for Scientific Research from the Japan Society for the Promotion of Science (JSPS; grant numbers #25253041, #17H01550 and 15H04764) and research funding from LSI Medience Corporation. We thank Edanz Group (www.edanzediting.com/ac) for editing a draft of this manuscript.

## Author Contributions

K.I. and T.U. designed the research and wrote the manuscript. S.U., K.G., S.M. and M.A carried out the experiments, obtained reagents and materials, and conducted data analysis. D.S. and N.A. contributed to network analysis. D.K, K.K. and T.U. contributed to flavonoid identification. All authors read and approved the final manuscript.

## Supporting information

**S1 Fig. Representative nanoparticle tracking analysis (NTA) of isolated m/lEVs.**

The distribution of the EV particle size from plasma (A) and urine (B) m/lEVs are indicated by density histogram.

**S2 Fig. Side-scatter setting with fluorescent particle and removal of non-specific binding reaction to IgG in flow cytometric observation.**

A. Development diagram of side-scatter and fluorescent intensity by using Fluorescent Particle Size Standard Kit.

(B and C) We selected mouse IgG negative group. It was subjected to the following characterization as plasma (B) and urine m/lEVs (C).

(D and E) Definition of Annexin V positive (Phosphatidylserine positive) and negative areas. After stained with fluorescent annexin V, EDTA was added at a final concentration of 10 mM and observed with a flow cytometer about plasma (D) and urine (E) fraction.

**S3 Fig. Gene enrichment analysis of proteins which are detected both plasma and urine m/lEVs in common.**

Top 20 clusters from Metascape pathway (http://metascape.org/) are shown.

**S4 Fig.Gating strategy for characterizing plasma m/lEVs by flow cytometry.**

A. After excluding the group which responded to normal mouse IgG (becomes positive), we shifted to the next step of characterization using the surface antigen from pre-set An5 + and - areas respectively.

B. Erythrocyte derived m/lEVs: After having gated a group of CD45 negative, we gated the group which was CD235a and CD59 double positive fraction.

C. Platelet derived m/lEVs: After having gated a group of CD5 and CD45 double negative, we gated the group which was CD41 and CD61 double positive fraction.

D. Macrophage/Monocyte/Granulocyte derived m/lEVs: After having gated a group of CD235a negative, we gated the group which was CD45 and CD15 double positive.

T cell/B cell derived m/lEVs: After having gated a group of CD235a negative, we gated the group which was CD45 and CD5 double positive fraction.

E. Endothelial cells derived m/lEVs: After having gated a group of CD146 positive (distinguished from the autofluorescence to rise from erythrocyte derived m/lEVs on the right), we gated the group which was CD105 positive fraction.

**S5 Fig. Gating strategy for characterizing urine m/lEVs by flow cytometry.**

After excluding the group which responded to normal mouse IgG (becomes positive), we shifted to the next step of characterization using the surface antigen from pre-set An5 + and - areas respectively.

MUC1 positive m/lEVs: After having gated a group of CD227 (MUC1) positive, we gated the group which was CD13 negative fraction.

Multiple peptidase (CD10, 13, 26) positive m/lEVs: After having gated a group of CD227 (MUC1) negative, we gated the group which was CD10, CD13 and CD26 triple positive fraction.

**S6 Fig. Quantification of DPPⅣ(CD26) enzymatic activity about plasma and urine m/lEVs.**

The distribution of the quantitative values of the three fractions (m/lEVs, Supernatant, Whole) is shown in a box-and-whisker plot (n=6 for human healthy plasma(A) and male urine(B), nmol/min/mL).

**S1 Table Plasma m/lEVs shotgun proteomics.**

**S2 Table Urine m/lEVs shotgun proteomics.**

**S3 Table The EV characterizing proteins which was detected by shotgun analysis.**

The list of the detected proteins which characterize EVs in MISEV2018. According to MISEV2018, we show proteins in line with Category 1, 2 and 4. XX = human gene names. XX* used for families of multiple proteins, for example for integrins: ITGA* indicates any integrin alpha chain.

P: The UniProt Accession number of the protein detected in plasma extraction compartment (Score).

U: The UniProt Accession number of the protein detected in urine extraction compartment (Score).

**S4 Table List of the proteins utilized for gene-enrichment analysis about plasma m/l EVs.**

We show the list which Log10(P-value) became lower than −20. Category: Classification of GO Terms, Term: GO term identifier, Description: Term name, Log10(P-value): −2 represents 0.01, the more negative the better, InTerm_InList: #Genes of Upload Hit List in this Term / #Genes of Genome in this Term, Symbols: List of Symbols of upload hits in this term.

**S5 Table List of the proteins utilized for gene-enrichment analysis about urine m/l EVs.**

We show the list which Log10(P-value) became lower than −20. Category: Classification of GO Terms, Term: GO term identifier, Description: Term name, Log10(P-value): −2 represents 0.01, the more negative the better, InTerm_InList: #Genes of Upload Hit List in this Term / #Genes of Genome in this Term, Symbols: List of Symbols of upload hits in this term.

